# Total neuroelectric brain activity derived from resting-state MEG is invariable across the adult lifespan

**DOI:** 10.1101/2024.06.21.600042

**Authors:** Mikhail Ustinin, Anna Boyko, Stanislav Rykunov

**Affiliations:** Keldysh Institute of Applied Mathematics, 125047 Russia; Russian Academy of Sciences Moscow, 125047 Russia

**Keywords:** magnetic encephalography, multichannel Fourier transform, mass solution of the inverse problem, current dipoles, age-related changes of the brain rhythms

## Abstract

Ageing of the human brain was studied using large array of experimental data. The magnetic encephalograms and magnetic resonance images of the head were obtained from the open archive CamCAN. Bad data were rejected, then functional tomograms were found - the spatial distribution of elementary spectral components. Physiological noise was eliminated by joint analysis of the functional tomograms and magnetic resonance images. By massively solving the inverse problem, multichannel spectra were transformed into time series of the power of elementary current dipoles. Age-related changes in the electrical power of various brain rhythms were examined. It was found that the summary electrical activity of the brain is constant throughout a person’s life. The electric power is redistributed during the lifetime: delta rhythm is diminishing, giving slow rise to all other rhythms.

## 1. Introduction

Age-related changes in brain activity are widely studied in modern neuroscience. Changes in cortical activity are considered (Rajah and D’Esposito 2005; Scally et al. 2018; Cespón et al. 2022) and also deeper compartments of the brain are investigated (Hinault, Baillet, and Courtney 2023; Shou et al. 2022; Mather and Nga 2013). Magnetic encephalography together with magnetic resonance imaging makes it possible to localize electrical sources with millimeter precision on a millisecond time scale (Hämäläinen et al. 1993; Llinás et al. 1999; Gómez et al. 2013; Baillet 2017). The main direction in the modern study of the brain ageing is the use of open databases, which contain the results of mass studies of various age cohorts (Van Essen et al. 2013; Shafto et al. 2014; Taylor et al. 2017; Niso et al. 2016). In this study we use the Cam-CAN (Shafto et al. 2014) repository as the biggest resource devoted to healthy ageing. It is used to study various features of ageing. In paper (Green et al. 2015) the question of cognitive variability across the adult lifespan was considered. Age-related changes in both rhythmic and arrhythmic signal strength in deeper brain regions were studied in (Hinault, Baillet, and Courtney 2023). The spatial correspondence between age effects on cortical thickness and those on functional networks were studied in (Stier, Braun, and Focke 2023). Recently, the method of the MEG-based functional tomography was proposed (Llinás and Ustinin 2014; Llinás et al. 2015; Ustinin, Boyko, and Rykunov 2023). It is based on detailed Fourier transform of the whole signal duration and massive solution of the inverse problem. This method presents all registered electrical activity of the body as a large set of elementary current dipoles. It was successively applied to study alpha rhythm, heart and skeletal muscles (Llinás et al. 2015, 2020; Rykunov, Boyko, and Ustinin 2023), in good agreement with anatomical data. Using this method, the magnetoencephalogram was divided into brain signal and physiological noise from the head (Llinás et al. 2022). The purpose of present work is to apply MEG-based functional tomography to the large set of data in order to study age-related changes of the total brain electrical activity and age dependence of various brain rhythms.

## 2. Materials and methods

### 2.1. Data acquisition

In this study we used MEG and MRI data downloaded from CamCAN (Shafto et al. 2014) data repository. The Cambridge Center for Aging and Neuroscience Research (Cam-CAN) Phase 2 cohort study archive is a large-scale (approximately 700 subjects), multimodal (MRI-magnetic resonance imaging, MEG-magnetic encephalography, cognitive study), cross-sectional population-based study of the mechanisms of cognitive aging spanning adult life expectancy (18-87 years). The dataset includes raw and pre-processed MRI, fMRI, MEG, and cognitive-behavioral data. The distribution of ages was chosen to be approximately equal, providing sufficient statistical power to test for differences both within and between age groups. Initial data quality control was performed by the Cam-CAN methodology team using semi-automated scripts.

#### 2.1.1. MRI acquisition

All MRI datasets were collected at a single location (MRC-CBSU, Cambridge, England) using a 3T Siemens TIM Trio with a 32-channel head coil. Participants were scanned in a one hour session. Before scanning, physiological measurements were taken and two behavioral experiments were performed. The scanner used memory foam cushions for comfort and to minimize head movements. Instructions and visual stimuli for functional tasks were back-projected onto a screen viewed through a mirror mounted on the head coil; auditory stimuli were presented through MR-compatible etymotic headphones; and manual responses were performed with the right hand using a custom-built MRI-compatible button unit. Cardiac data were recorded using a photoplethysmograph/pulse oximeter on the left index finger at 50 Hz.

#### 2.1.2. MEG acquisition

All MEG data sets were collected at a single location (MRC-CBSU, Cambridge, England) using a 306-channel VectorView MEG system (Elekta Neuromag, Helsinki) consisting of 102 magnetometers and 204 orthogonal planar gradientometers located in a magnetically shielded room (MSR). Data were sampled at 1 kHz with a 0.03 Hz upper-pass filter. Recordings were made in a sitting position. Head position in the MEG helmet was continuously assessed using four head position indicator (HPI) coils to allow autonomous correction of head motion. Two pairs of bipolar electrodes were used to record vertical and horizontal electrooculogram (VEOG, HEOG) signals to monitor blinks and eye movements, and one pair of bipolar electrodes recorded an electrocardiogram (ECG) signal to monitor heart rate-related artifacts. Instructions and visual stimuli were projected onto a screen through an opening in the front wall of a magnetically shielded room; auditory stimuli were presented through etymotic tubes; responses were made through a custom-made push-button box with fiber-optic wires. MEG data were collected at rest, during sensorimotor and audiovisual (passive) tasks. During the resting state recording, participants sat motionless with eyes closed for at least 8 minutes and 40 seconds.

### 2.2. Data preprocessing

#### 2.2.1. MEG data

To suppress external noises, we applied Maxwell filter (Taulu and Kajola 2005; Taulu and Simola 2006) routine from MNE (Gramfort et al. 2013) package to the raw MEG data. Time series of magnetometer channels were selected for further analysis. On the next step time series were trimmed to 300 seconds length, starting from 100th second of recording. Using the FFTW library (Frigo and Johnson 2005), multichannel spectra were computed over the entire length (300 sec) of the time series. Data structures describing the spatial configuration of the magnetic encephalograph and its position in space relative to the subject’s head were constructed using the FieldTrip (Oostenveld et al. 2011) package.

#### 2.2.2. MRI data

MRI images were realigned to CTF-like coordinate system^1^ with 2mm spatial resolution. To construct an annotated brain map, we segmented magnetic resonance images of the human head using a pre-trained UNesT neural network (Huo et al. 2019; Zhang et al. 2022). The weights for this network were taken from the Project MONAI (Cardoso et al. 2022) repository. The ANTsPy (Tustison et al. 2021) package was used for translation into MNI (Montreal Neurological Institute coordinate system, (Fonov et al. 2009)) space and vice versa. For the purposes of this study, we combined into a single region of interest the many segmentation regions that together make up the whole brain for every subject’s MRI.

### 2.3. MEG-based functional tomography

Let us consider the theory and experimental justification of the MEG-based functional tomography, as they were set out in the works (Llinás and Ustinin 2014; Llinás et al. 2015; Ustinin, Boyko, and Rykunov 2023). The encephalograph simultaneously records the values of magnetic field induction in *K* channels during time *T* producing a set of experimental vectors {***B***_***k***_}, *k =* 1, …, *K*. The field is recorded at *H* points in time with a constant step *T/H*. As a result of the measurements, we obtain a two-dimensional data array ***B***_***hk***_, where *h =* 0, …, *H* − 1, *k =* 1, …, *K*. Multichannel discrete Fourier transform can be written as:

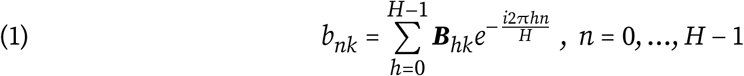

where *b*_*nk*_ - complex Fourier amplitude for frequency ν_*n*_ in a channel with a number *k*, and 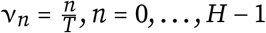. All spectra are computed for the full measurement time *T*, which reveals the detailed frequency structure of the system. Frequency resolution is equal to 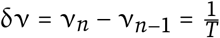, thus, the frequency resolution is determined by the registration time. The phase and amplitude of the *n*-th component of the Fourier series are written as: φ_*nk*_ = *atan*2(*Imb*_*nk*_, *Reb*_*nk*_), 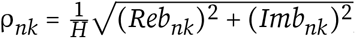, where *Reb*_*nk*_ – real part of *b*_*nk*_, *Imb*_*nk*_ – imaginary part.

Knowing the multichannel spectrum, the inverse Fourier transform can be performed on all channels:

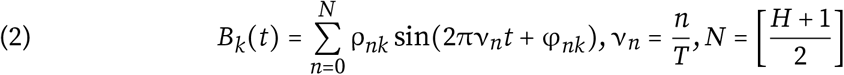

In order to study the detailed frequency structure of the brain, we reconstruct the multi-channel signal at each frequency and analyze the obtained time series. Reconstructed multichannel signal of frequency ν_*n*_ in all channels:

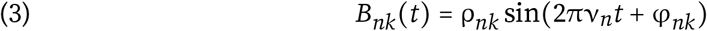

where 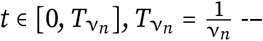 period of that frequency.

Figure 1B shows a map of the magnetic field at one frequency. It can be noted, that such field configuration can be accurately described by a single equivalent current dipole (Sarvas 1987):

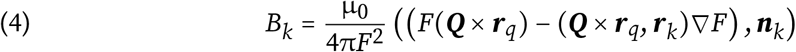

where 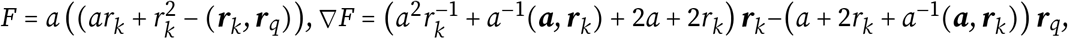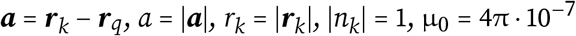

**FIGURE 1.**
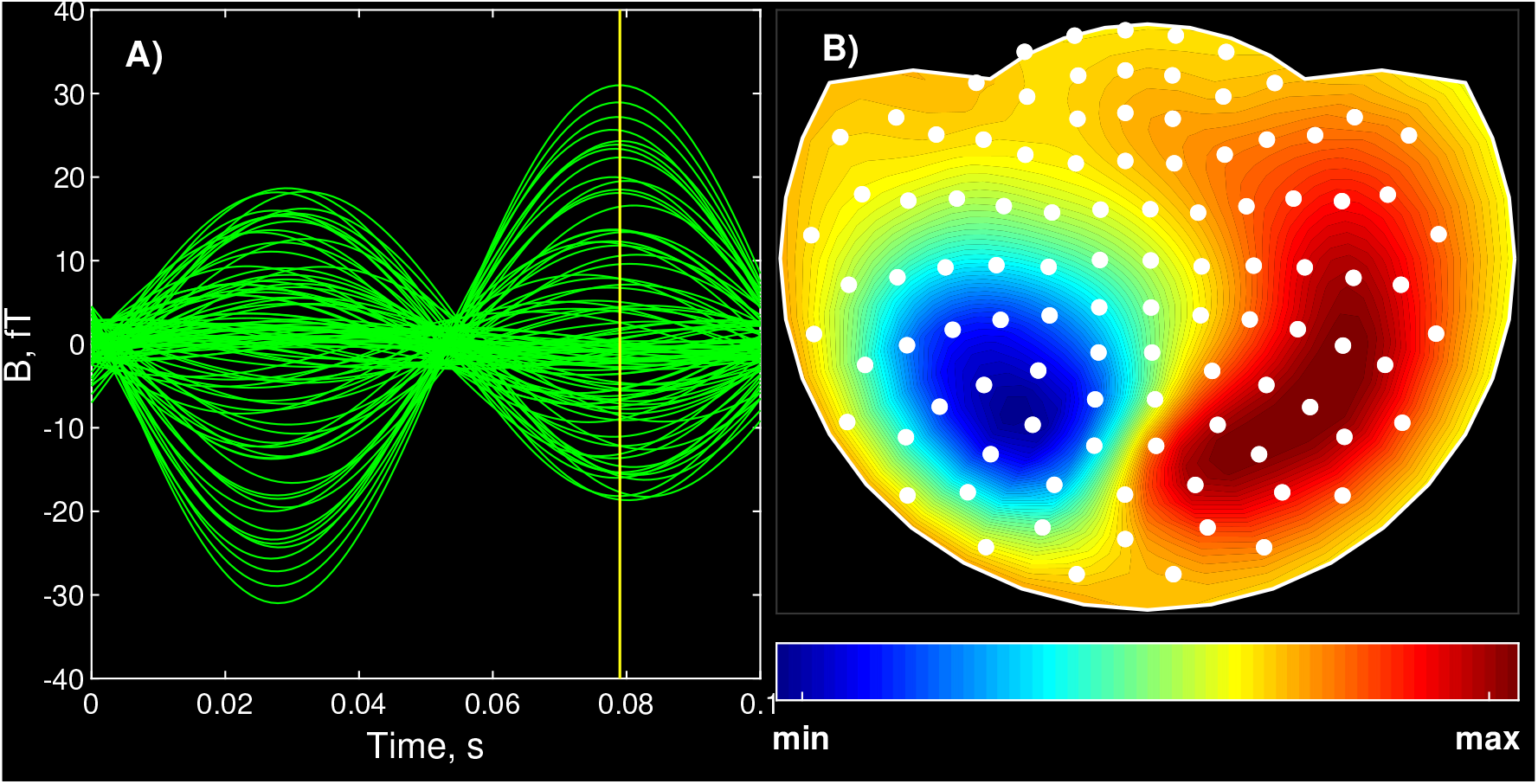
A) Reconstructed multichannel time-series at one frequency(ν_*n*_ = 9.79*Hz*). Yellow line marks moment of maximum; B) Normalized map of the magnetic field taken at point of maximum. White outline is equidistant projection of MEG helmet on a plane, white circles marks channels positions.

Considering Figure 1A, one can see very clean multichannel signal with extremely high coherence. From Figure 1B follows, that the source of this signal is a single equivalent current dipole. So, simply performing direct and inverse Fourier transform of the experimental magnetoencephalogram, we can look inside the brain at the particular frequency. Such phenomena were observed in (Llinás and Ustinin 2014) and became an experimental basis of the model in which one frequency component is described by one current dipole. We’ve found, that the magnitude ratios between channels remain constant, only the scale changes. Thus formula (3) can be written as:

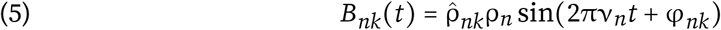

where 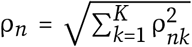 amplitude, 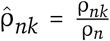 value of normalized pattern 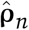 of this oscillation in *k*-th channel. In multichannel measurements, the space is determined by the channel arrangement. The use of normalized patterns makes it possible to determine the spatial structure of the signal sources by the solution of the inverse problem, and this structure remains constant over the entire oscillation time. The time dependence of the field is determined by the function ρ_*n*_ sin(2πν_*n*_*t* + Φ_*n*_), which is common for all channels, i.e., this source oscillates as a single unit at frequency ν_*n*_.

The theoretical foundations for the reconstruction of static functional entities (neural circuits, or sources) were outlined in (Llinás and Ustinin 2014; Llinás et al. 2015). This reconstruction is based on detailed frequency analysis and the selection of frequency components with similar patterns. Algorithm for this method can be written as:

1. Fourier transform of the input multichannel signal.
2. Inverse Fourier transform – reconstruction of the signal at each frequency.
3. Calculation of phase, normalized pattern and its amplitude for each frequency component.
4. Solution of the inverse problem for all of the components.

To calculate phases, normalized patterns of experimental signal components and their amplitudes we use the following procedure for each frequency:

1. Reconstruct a multichannel signal at a single frequency for a single period of that frequency.
2. Find a maximum of that reconstructed signal. For this purpose, we calculate the sum of squares of amplitudes in all channels at each point in time.
3. At the point of maximum, we take a “slice” of the multichannel signal, representing a vector of length k, and calculate the norm of this vector. This normalized vector is a pattern 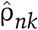 of selected frequency.
4. Phase estimation for this frequency:

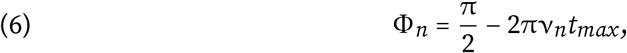

where *t*_*max*_ is moment of maximum, found on step 2.

Each elementary oscillation is characterized by frequency ν_*n*_, phase Φ_*n*_, amplitude ρ_*n*_, normalized pattern 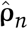, and its source is a functional entity with a permanent spatial structure. The functional tomography method reconstructs the structure of the system by analyzing a set of normalized patterns 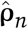. A functional tomogram shows a three-dimensional map of the distribution of energies produced by sources located at a given position in space. To construct a functional tomogram, the investigated region of space is divided into *N*_*x*_ ×*N*_*y*_ ×*N*_*z*_ elementary cubic cells with centers at ***r***_*ijs*_. The length of the cube edge is chosen according to the desired accuracy, in this study it was 2 mm. In each cell set of test dipoles ***Q***_*ijsl*_ with direction ***L*** is constructed. The magnetic induction generated by the test dipole ***Q***_*ijsl*_, located at the point ***r***_*ijs*_ is recorded by a sensor with the number *k*, located at the point with coordinates ***r***_*k*_ and having the direction ***n***_*k*_, *k*-th component 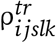 of the test pattern 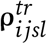 is calculated using the model of the current dipole in a spherical conductor (Sarvas 1987):

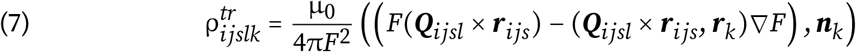

where 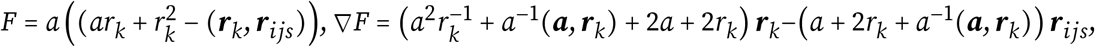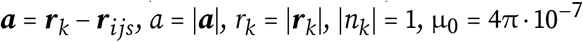 The normalized pattern is calculated as:

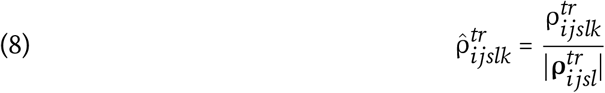

where 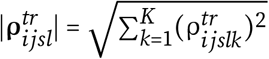. All test dipoles located at the point ***r***_*ijs*_, lie within the same plane orthogonal to ***r***_*ijs*_, since the result of the vector product ***Q***_*ijsl*_ × ***r***_*ijs*_ is non-zero only for such dipoles. The test dipoles cover the circle in *L*_*max*_ directions in 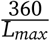 degree increments, *L*_*max*_ = 72 was used in this study. For each of the dipoles, a set of normalized patterns is computed using formula (7):

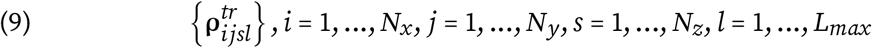

For each of the normalized experimental patterns 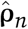 the following function is computed to determine the difference between that pattern and one of the test patterns:

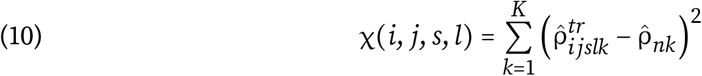

where 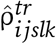 – *k* -th component of the normalized trial pattern 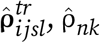 – *k* -th component of the normalized experimental pattern 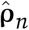, *k* – channel number. Position and direction of the source corresponding to the pattern 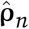, is determined by the numbers (*I, J, S, L)*, corresponding to the minimum of the function χ (*i, j, s, l)* on variables *i* 1, …, *N*_*x*_ ; *j* 1, …, *N*_*y*_; *s* 1, …, *N*_*z*_ ; *l* 1, …, *L*_*max*_. The minimum of this function is found by the method of exhaustive search, i.e., by selecting the smallest of several million values of the function χ for each pattern 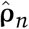. This procedure determines the position ***r***_*IJS*_ – solution of the inverse problem for pattern 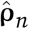, without spatial filtering of channels and without introduction of weight functions. In this study, we propose a method for direct estimation of the amplitudes of the dipole sources producing the measured magnetic field:

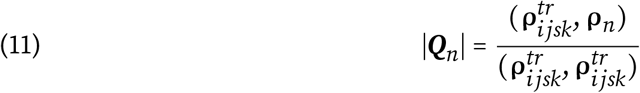

Repeating this procedure for all normalized patterns 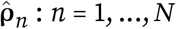, it is possible to find position, direction and amplitude of the source dipole for all frequency components of the initial experimental signal, thus solving the inverse problem of magnetic encephalography. After solution of the inverse problem, a functional tomogram, a three-dimensional distribution of sources in space, is obtained. Each cell of space corresponds to a set of sources characterized by their frequencies, dipole moments and their spectral powers.

### 2.4. Data processing and analysis

Functional tomograms were computed for all 595 experimental data sets with the following parameters: time series duration 300 sec, frequency bandwidth 0.3 − 100*Hz*, spatial resolution 2*mm*, number of directions 72. On the next step of analysis each functional tomogram was separated into two parts: one containing sources localized in the brain, other containing all other physiological sources. Such separation was achieved by combining functional tomogram and whole brain mask, acquired on MRI data preprocessing stage. Results of separation are shown on Figure 2.

**FIGURE 2.**
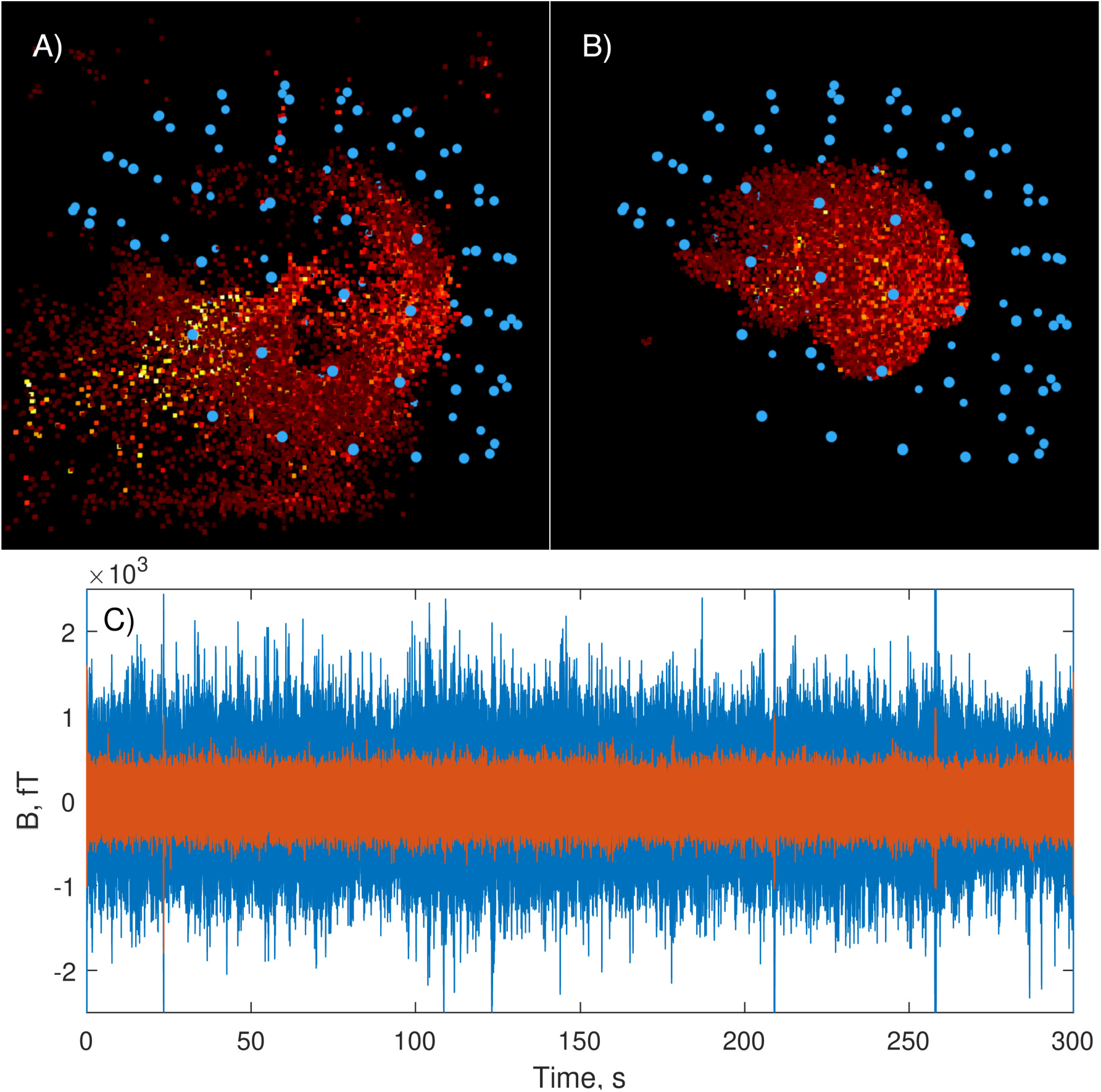
A)functional tomogram of non-brain sources; B)Functional tomogram of brain sources; C) Reconstructed time series for brain (blue) and non-brain(orange) sources

After that time series were reconstructed for all brain functional tomograms and total magnetic energies were calculated. The outliers were found by median filtering. The energies of these outliers differed from the median values by several orders of magnitude. These experimental data were discarded from further analysis. This procedure left 501 experimental sets. The remaining sets were then grouped by age. The number of experiments in each age group is presented in Table 1. For each group powers were calculated in conventional frequency bands (0.3-4 Hz – delta; 4-8 Hz – theta; 8-13 Hz – alpha; 13-21 Hz – beta 1; 21-30 Hz – beta 2; 30-98 Hz – gamma) and averaged.

**TABLE 1.** Distribution of experiments by age groups.

| age | 18-24 | 25-29 | 30-34 | 35-39 | 40-44 | 45-49 | 50-54 | 55-59 | 60-64 | 65-69 | 70-74 | 75-79 | 80-88 |
| --- | --- | --- | --- | --- | --- | --- | --- | --- | --- | --- | --- | --- | --- |
| number of experiments | 16 | 38 | 33 | 38 | 41 | 45 | 39 | 37 | 36 | 45 | 32 | 48 | 53 |

## 1. Results

### 3.1. MEG spectral power

In this study, the MEG spectral power is understood as the sum of the spectral powers of all frequencies falling within the range of a given rhythm per unit time. Figures 3-5 illustrate age dependence of the MEG spectral power. The following frequency bands were considered: 0.3-98 Hz - total spectral power; 4-98 Hz – total spectral power excluding delta rhythm; 0.3-4 Hz – delta rhythm; 4-8 Hz – theta rhythm; 8-13 Hz – alpha rhythm; 13-21 Hz – beta 1 rhythm; 21-30 Hz – beta 2 rhythm; 30-98 Hz – gamma rhythm.

**FIGURE 3.**
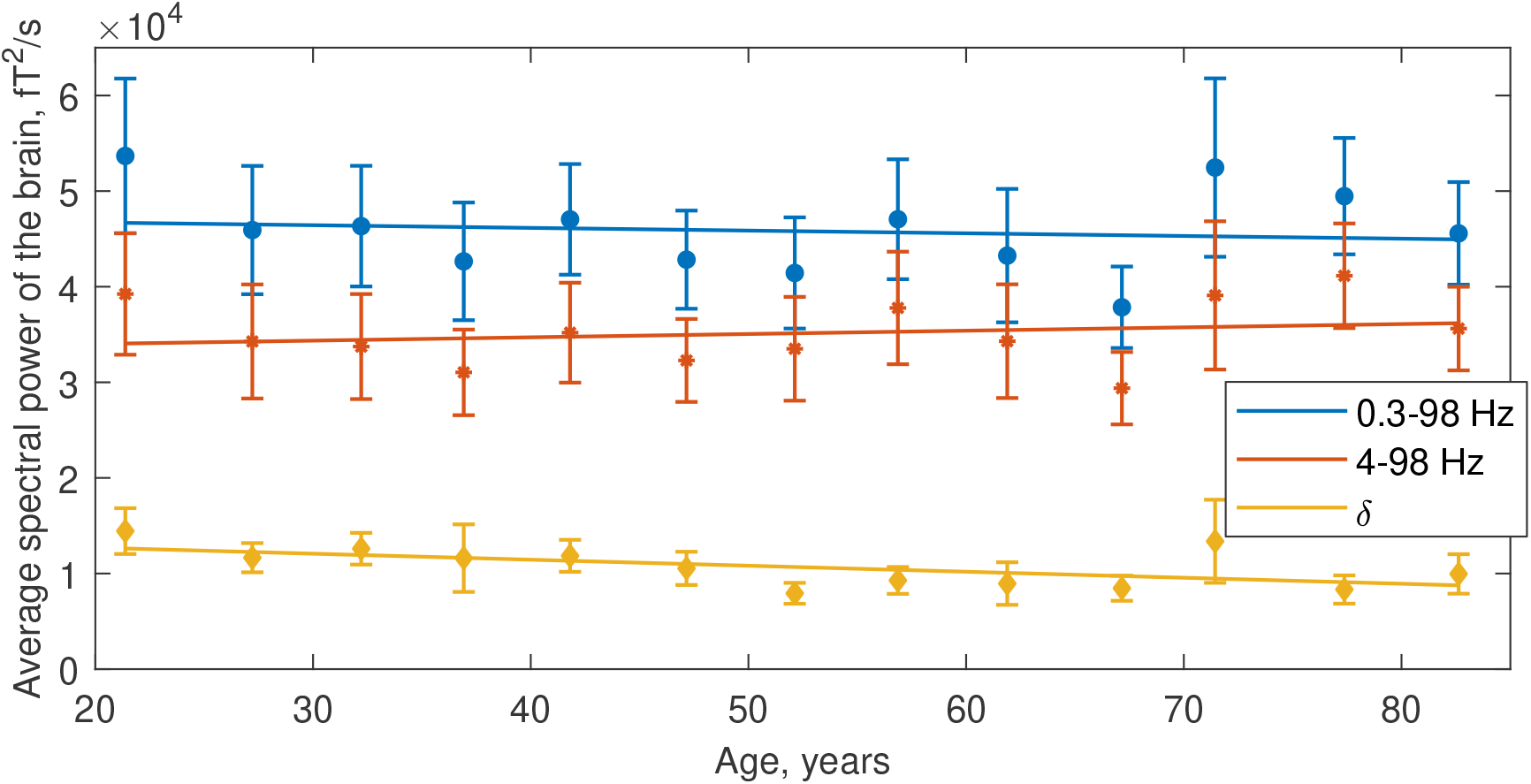
Dependence of the average MEG spectral power on age. Blue dots and line – total spectral power in the frequency band 0.3 − 98*Hz*. Orange dots and line – spectral power in the frequency band 4 − 98*Hz* (without delta rhythm). Yellow dots and line – spectral power in the delta frequency band 0.3 − 4*Hz*

As can be seen in Figure 3 (blue dots and line), total spectral power in the frequency band 0.3-98 Hz shows some downward trend, although the linear decay is not significant. Age dependence of power in the same band excluding delta rhythm (red dots and line) shows some upward trend, while the linear growth is also not significant.

Figure 4 shows a comparison of all rhythms on a single scale. In Figure 5 age dependences of rhythms are shown separately in their own scale to show more details in low-amplitude rhythms. As can be seen in Figure 4, alpha rhythm (yellow dots and line) makes the largest contribution to the MEG total spectral power and decreases over time. Delta rhythm (blue dots and line) also makes a significant contribution and decreases with age. The beta 1 rhythm is definitely increasing (purple dots and line). Theta rhythm (orange) is more or less constant through the lifespan. As can be seen in Figure 5, beta 2 (green) and gamma (light blue) are small and slowly rising. It can be concluded that the decrease in the MEG spectral power of the alpha and delta rhythms is compensated by the increase in sum of all other rhythms.

**FIGURE 4.**
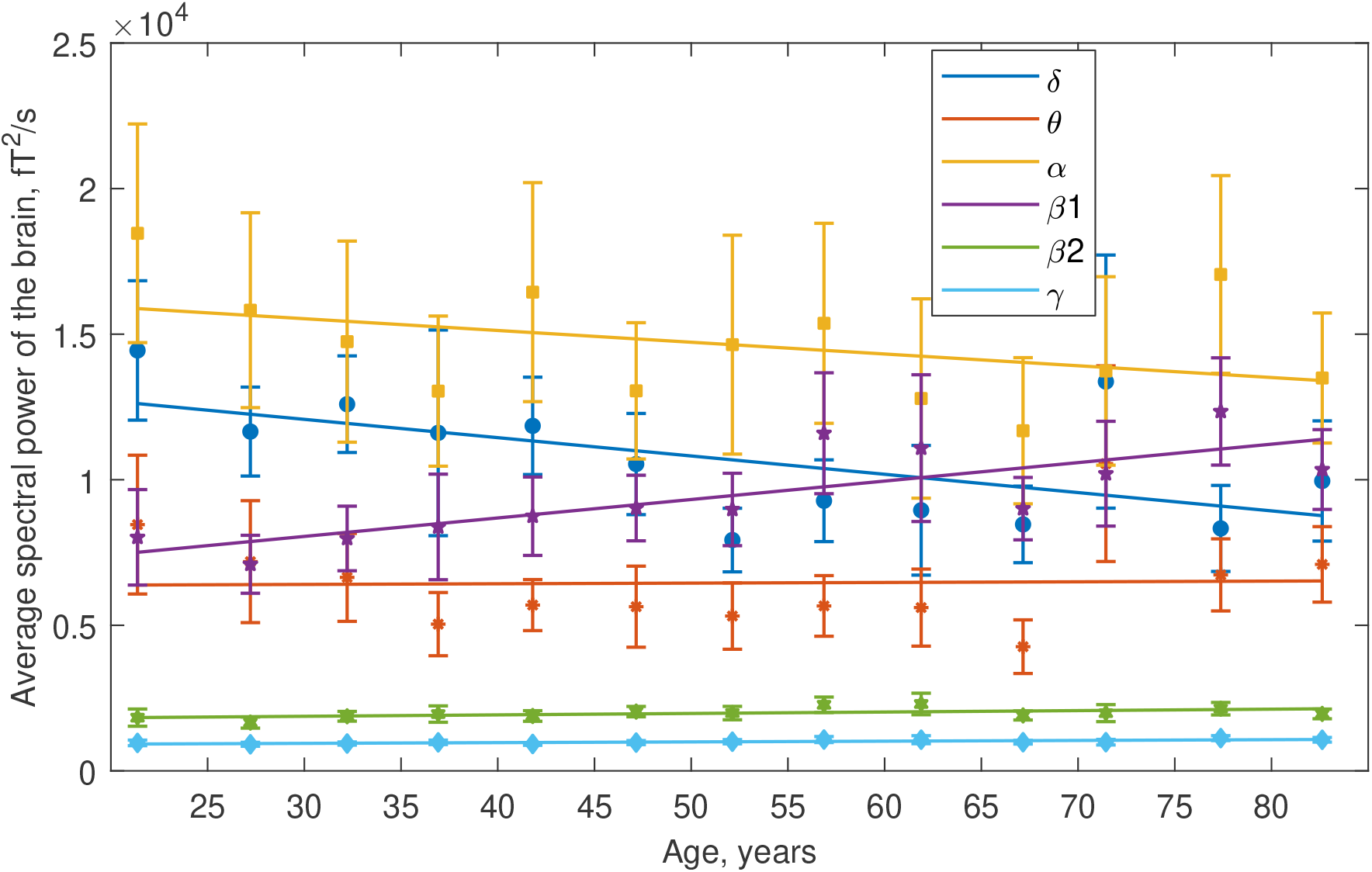
Dependence of average MEG spectral power on age, all brain rhythms. Colors corresponding to different rhythms are indicated in the subpanel.

**FIGURE 5.**
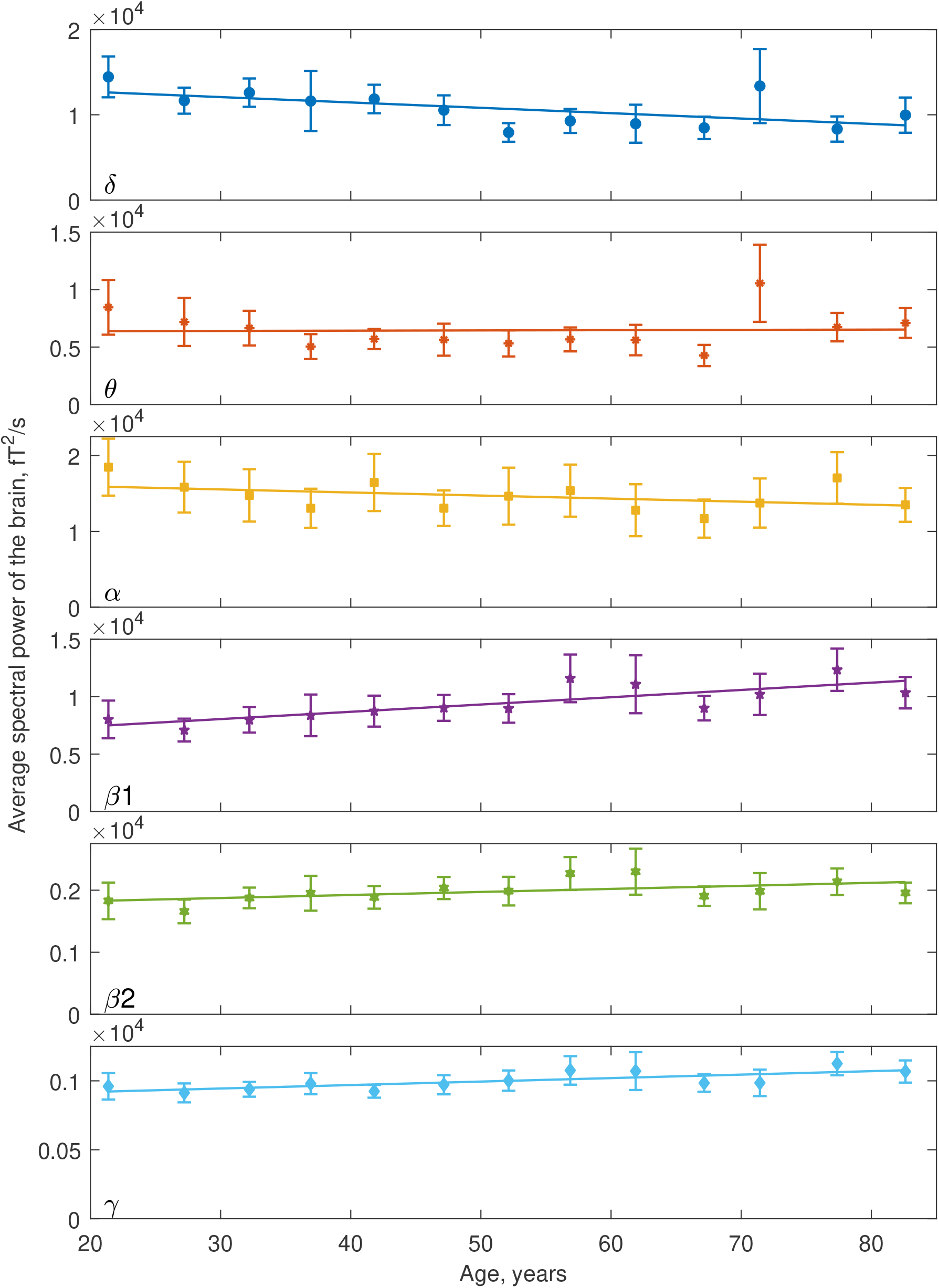
Dependence of average MEG spectral power on age, all brain rhythms.

### 3.2 The electric power of brain sources

The electric power of the elementary dipolar source can be defined as total energy, produced by this source per unit time:

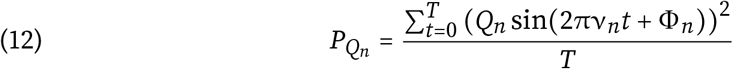

Electric power of the brain in particular frequency band is a sum of powers of all sources located in the brain, whose frequency lies in this band.

By mass solving of the inverse problem, we calculated the time dependence of the summary electric power of elementary current dipoles reflecting whole brain activity. Figures 6–8 illustrate age dependence of the brain electric power. Typical number of the brain dipoles in our model is approximately 30 thousand for frequency band 0.3-98 Hz. The following frequency bands were considered: 0.3-98 Hz - total electrical power; 4-98 Hz – total electrical power excluding delta rhythm; 0.3-4 Hz – delta rhythm; 4-8 Hz – theta rhythm; 8-13 Hz – alpha rhythm; 13-21 Hz – beta 1 rhythm; 21-30 Hz – beta 2 rhythm; 30-98 Hz – gamma rhythm.

**FIGURE 6.**
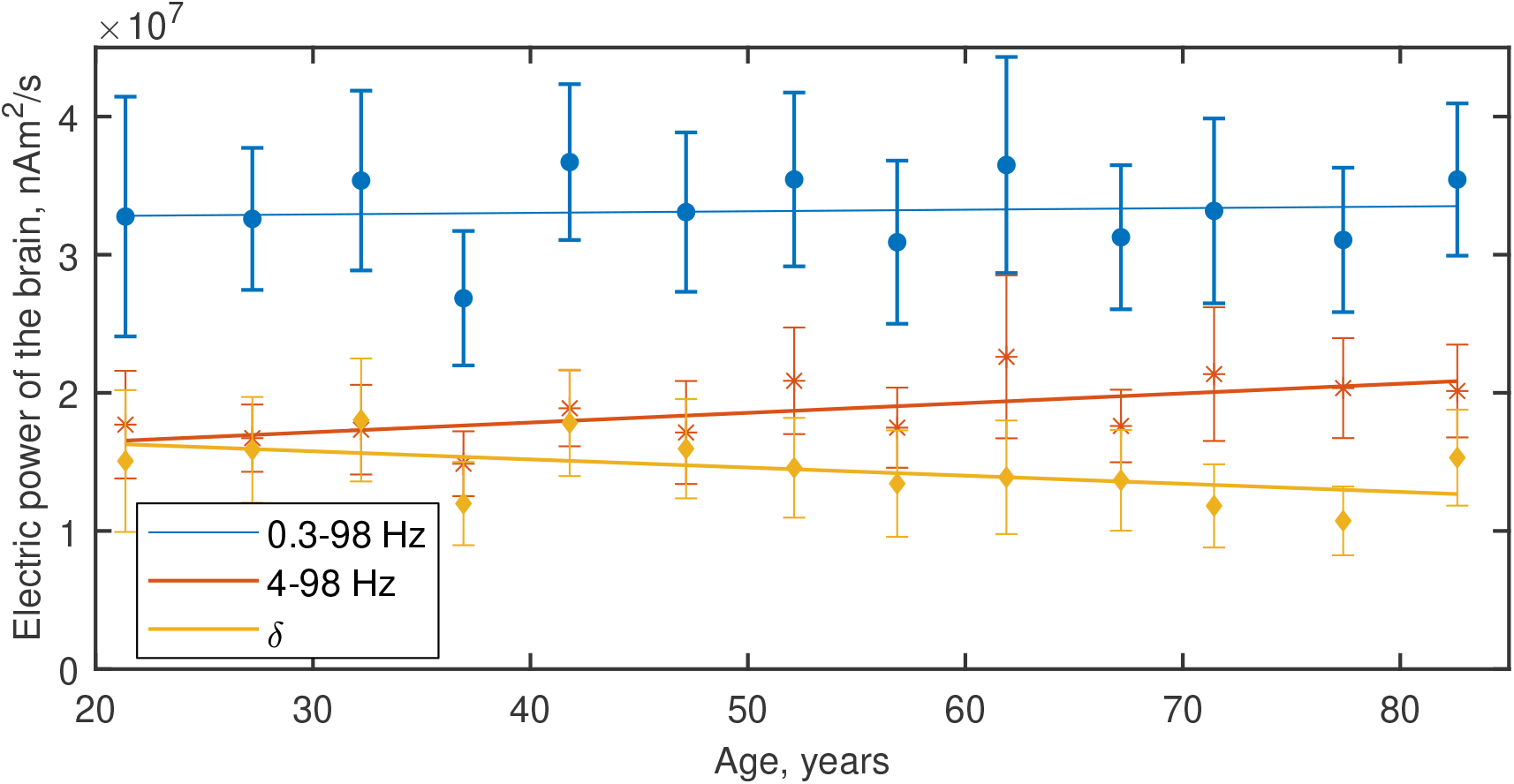
Dependence of the brain average electric power on age. Blue dots and line — total electrical power in the frequency band 0.3-98 Hz. Red dots and line — electric power in the frequency band 4-98 Hz (excluding delta rhythm). Yellow dots and line – electric power in the delta frequency band 0.3 − 4*Hz*

As can be seen in Figure 6 (blue dots and line), total electrical power in the frequency band 0.3-98 Hz (blue dots and line), is constant through the lifespan, with minor stochastic oscillations between age cohorts. Age dependence of electric power in the delta rhythm band (yellow dots and line) shows downward trend, while the sum of all other rhythms (orange dots and line) increases at the same rate.

Figure 7 shows a comparison of all rhythms on a single scale. In Figure 8 age dependences of rhythms are shown separately in their own scale to show more details in low-amplitude rhythms. Considering the Figure 7, one can see, that unlike the MEG spectral power (see Figure 3), delta rhythm (blue dots and line) makes the largest contribution to the brain electrical power and decreases with age. From the Figures 7 and 8 follows, that electrical powers of the alpha rhythm (yellow dots and line), beta 1 rhythm (purple dots and line), theta rhythm (orange dots and line), beta 2 rhythm (green dots and line) and gamma rhythm (light blue dots and line) grow with age. It can be concluded that the decrease in the electrical power of the delta rhythm is compensated by the summary growth in the sum of all other rhythms.

**FIGURE 7.**
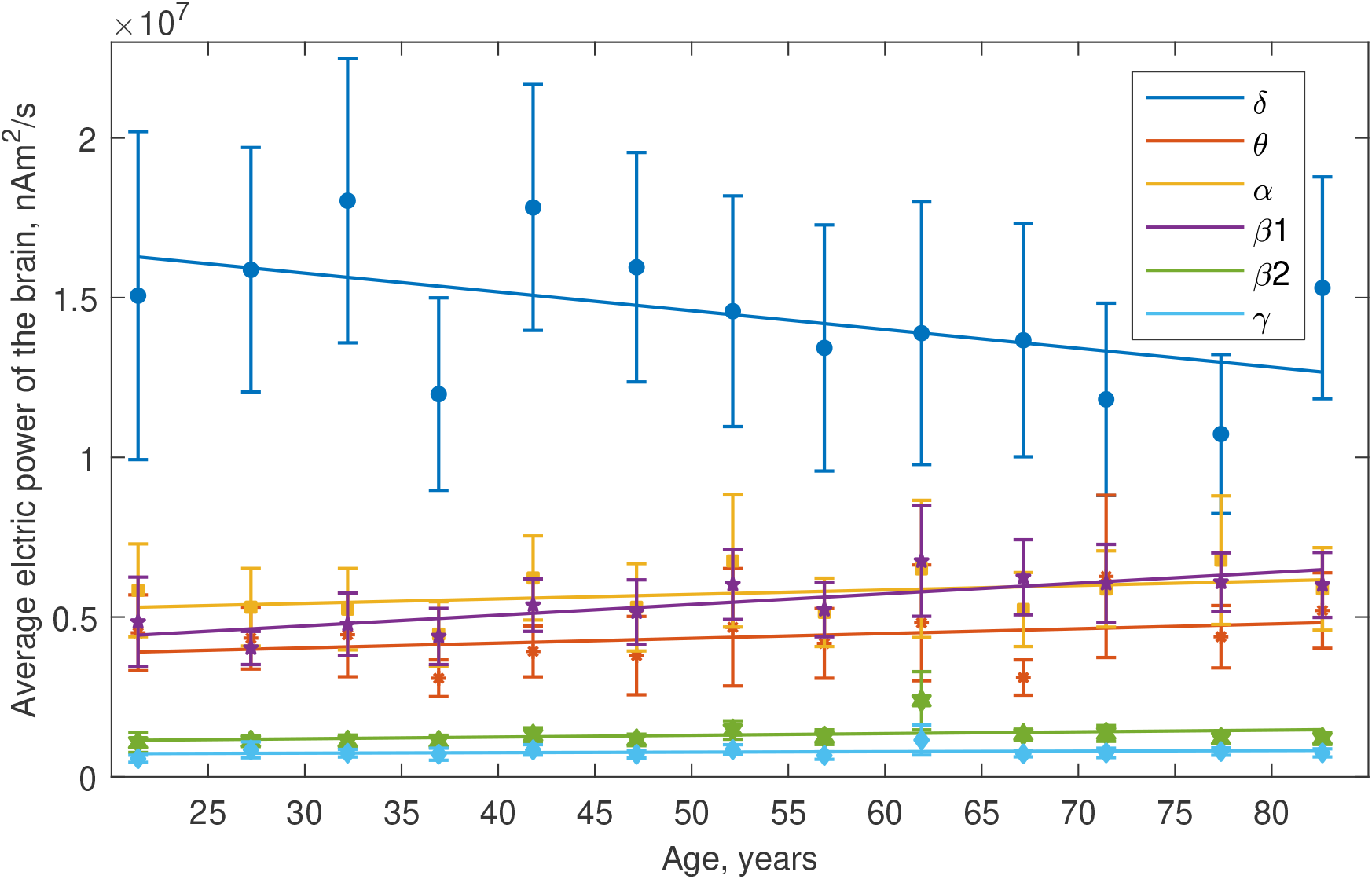
Dependence of average electrical power on age, all brain rhythms. Colors corresponding to different rhythms are indicated in the subpanel.

**FIGURE 8.**
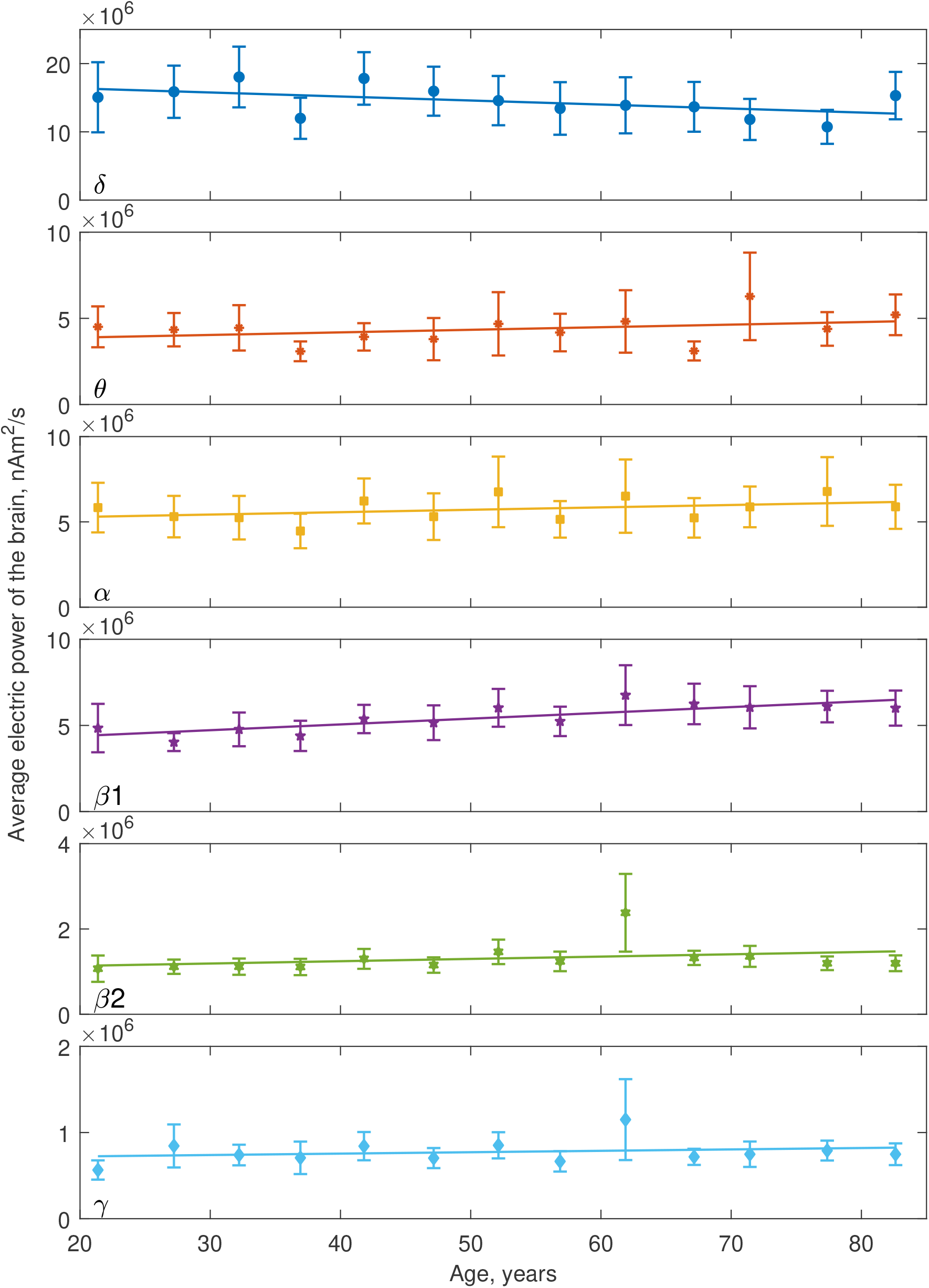
Dependence of average electric power on age, all brain rhythms.

## 4. Discussion

Authors of the work (Stier, Braun, and Focke 2023) probed lifespan differences in power and connectivity. Let us compare their results for spectral power with Figure 4. MEG power in (Stier, Braun, and Focke 2023) was computed for six conventional frequency bands based on fast Fourier spectral analysis. Time dependencies for delta, theta, beta and gamma rhythms, are similar to our results, while alpha rhythm differs. In (Stier, Braun, and Focke 2023) alpha rhythm is slowly growing, at Figure 4 it is slowly falling. Still, if we look at the time dependence of the electric power of the alfa rhythm sources, it is definitely growing. In paper (Hoshi and Shigihara 2020) the inverse problem was solved using the sLORETA-like algorithm (Pascual-Marqui 2002). The regional oscillatory power for several frequency bands was estimated in order to study the effects of healthy aging and gender asymmetricity on the regional resting-state brain activity.

We suppose, that true time dependencies are revealed after the solution of inverse problem, because moving of the sources with age can change MEG data, even if the source power is constant. Comparing Figures 4 and 7, one can note very different roles of the spectral and electrical powers of the delta rhythm. Spectral delta-power is less than alpha and comparable to beta 1. Electrical delta-power is dominating for all ages, which can be explained by the deep position of its sources, including thalamic area. Invariance of the total electric power through the life span is the result of the compensation of the delta electric power falling by the growth of the summary theta, alpha and beta 1 electric power. More detailed study of the spatial distribution of electrical sources and underlying neurological mechanisms can be performed using approach proposed in present article.

## 5. Conclusion

In this work we applied the newly developed method of MEG-based functional tomography to large CamCAN archive studying the ageing of human brain. It was found, that time dependencies of the brain rhythms spectral powers differ from those for electric powers revealed by the mass solution of inverse problem. Two main conclusions can be made. First, the total electric power of the brain remains constant through the human lifespan. Second, during the lifetime electric power is redistributed: electric power of the delta rhythm is diminishing, giving slow rise to the theta, alpha and beta 1 rhythms.

## Footnotes

1 https://www.fieldtriptoolbox.org/faq/coordsys/

